# Mycelial nutrient transfer promotes bacterial co-metabolic organochlorine pesticide degradation in nutrient-deprived environments

**DOI:** 10.1101/2022.10.10.511588

**Authors:** Nelson Khan, Edward Muge, Francis J. Mulaa, Benson Wamalwa, Martin von Bergen, Nico Jehmlich, Lukas Y. Wick

**Author notes:** Corresponding author: Lukas Y. Wick, Mailing Address.

## Abstract

Biotransformation of soil organochlorine pesticides (OCP) is often impeded by a lack of nutrients relevant for bacterial growth and/or co-metabolic OCP biotransformation. By providing space-filling mycelia, fungi promote contaminant biodegradation by facilitating bacterial dispersal and the mobilization and release of nutrients in the mycosphere. We here tested whether mycelial nutrient transfer from nutrient-rich to nutrient-deprived areas facilitates bacterial OCP degradation in a nutrient-deficient habitat. The legacy pesticide hexachlorocyclohexane (HCH), a non-HCH-degrading fungus (*Fusarium equiseti* K3) and a co-metabolically HCH-degrading bacterium (*Sphingomonas* sp. S8) isolated from the same HCH-contaminated soil were used in spatially structured model ecosystems. Using ^13^C-labelled fungal biomass and protein-based stable isotope probing (protein-SIP), we traced the incorporation of ^13^C fungal metabolites into bacterial proteins while simultaneously determining the biotransformation of the HCH isomers. The relative isotope abundance (RIA, 7.1 – 14.2%), labeling ratio (LR, 0.13 – 0.35), and the shape of isotopic mass distribution profiles of bacterial peptides indicated the transfer of ^13^C-labeled fungal metabolites into bacterial proteins. Distinct ^13^C incorporation into the haloalkane dehalogenase (linB) and 2,5-dichloro-2,5-cyclohexadiene-1,4-diol dehydrogenase (LinC), as key enzymes in metabolic HCH degradation, underpin the role of mycelial nutrient transport and fungal-bacterial interactions for co-metabolic bacterial HCH degradation in heterogeneous habitats. Nutrient uptake from mycelia increased HCH removal by twofold as compared to bacterial monocultures. Fungal-bacterial interactions hence may play an important role in the co-metabolic biotransformation of OCP or recalcitrant micropollutants (MPs).

## Introduction

Despite its ban or restricted use in many parts of the world, the organochloride pesticide hexachlorocyclohexane (HCH) continues to pose a serious environmental risk (1,2) due to its toxicity and environmental persistence (1,3–5). Commercial production of HCH typically results in a mixture of four major isomers (*α*-HCH (60 – 70%), *β*-HCH (5 – 12%), *γ*-HCH (10 – 12%), *δ*-HCH (6 – 10%)) (3,6), of which only *γ*-HCH has insecticidal activity while the other isomer are discarded as HCH-muck, thereby generating stockpiles of persistent toxic waste (3,6–8). Microorganisms including fungi (5,9,10) and bacteria (7,11,12) are able to degrade HCH isomers. However, an elaborate degradation pathway and the respective catabolic enzymes for complete mineralization of HCH have only been reported for bacterial members of the family sphingomonadaceae under ideal laboratory conditions (3,7,11,12). Microbial degradation in heterogeneous environments though emerges from manifold biotic and abiotic factors including redox conditions, moisture, and – in cases of co-metabolic degradation – the presence and spatial availability of auxiliary organic growth substrates to potential degraders (13–15). Contrary to bacteria, mycelial fungi are highly adapted to heterogeneous habitats. Due to their space-filling mycelial structures, they translocate resources within their networks from nutrient-rich to nutrient-deficient patches (16,17).

Fungi and bacteria thereby often share the same microhabitat and contribute to nutrient cycling and other biogeochemical processes (17–19), such as the biotransformation of organic contaminants (14,20,21). In the mycosphere (i.e. the microhabitat that surrounds fungal hyphae and mycelia), bacteria and fungi may compete for the same substrates and, depending on the organisms and habitat conditions, their interactions may be either ecologically neutral, competitive, or cooperative (22,23) and range from apparently random physical to specific commensal or symbiotic associations (19). Mycelia e.g. have been shown to actively transport polycyclic aromatic hydrocarbons (PAH) thus improving their bioavailability to bacterial degradation in unsaturated environments (24–26). They may further serve as transport vectors for bacteria and phage-bacteria couples (27), thus improving the access of bacteria to otherwise poorly available nutrients and contaminants(20,28). Fungi are also known to shape bacterial community structures surrounding the hyphosphere by secreting carbonaceous compounds that can be utilized by fungus-associated bacteria (17,25,29,30). They thereby exude carbon substrates such as phenols, citrate, or oxalate that may serve as auxiliary carbon sources for pollutant-degrading bacteria (21).

Co-metabolic degradation offers the benefit of contaminant removal even at trace quantities since the degrader is not reliant on the contaminant for carbon or energy source (15,31). However, the role of fungal resource transfer to bacteria for improved bacterial activity and co-metabolic contaminant degradation in contaminated oligotrophic microhabitats remains vastly underexplored. In microbial ecology, stable isotope probing of proteins (protein-SIP) has become a valuable technique that allows the identification of key metabolic actors and functional interactions in microbial communities (32–34). Protein-SIP combines meta-proteomics based on mass spectrometry and stable isotope probing techniques to sensitively detect the assimilation of labeled target substrate (such as ^13^C) at peptide level (32,34). The information gleaned from mass spectrometry data includes relative isotope abundance (RIA), labeling ratio (LR), and the shape of the isotopic distribution (32). The RIA describes the percentage of the heavy isotope (i.e., ^13^C) atoms in a peptide, which provides information on the incorporation of the labeled substrate into biomass. The LR describes the ratio of labeled to unlabeled peptides which relate to protein turnover following the addition of the labeled substrate (32,35).

In the present study, we used spatially fragmented laboratory model ecosystems to test the hypothesis whether mycelial nutrient transfer from eutrophic habitats promotes co-metabolic degradation of HCH by bacteria in oligotrophic contaminated environments. A non-HCH-degrading soil fungus *Fusarium equiseti* K3 and a co-metabolically HCH degrading bacterium *Sphingobium* sp. S8 isolated from the same HCH contaminated soil were used. Strain S8 was placed onto a nutrient-free HCH-treated agar patch that was separated from the fungal inoculum by an air gap. ^13^C-labeled mycelia of strain K3 were allowed to overgrow the bacterial region. Using a combination of metaproteomics and stable isotope labeling and analyses (34), we traced the incorporation of ^13^C of fungal origin into bacterial proteins responsible for both housekeeping and co-metabolic HCH degradation respectively, and simultaneously followed by chemical analytics. Our results demonstrate that mycelial fungi facilitate bacterial degradation of contaminants in nutrient-deficient environments by providing C-substrates to bacteria, thus enhancing their biotransformation capacity.

## Materials and Methods

### Chemicals

Hexane (LiChrosolv grade), Acetone (99.8%), and γ-HCH (97%) were either purchased from Sigma (Munich, Germany) or Merck (Darmstadt, Germany) and were of analytical grade. ^13^C-glucose (D-glucose^13^C_6_, 99%) was obtained from Cambridge Isotope Laboratories, Inc, (USA) The analytical standard containing a mixture of the HCH isomers, *α*: *β*: *γ*: *δ* = 1:1:1:1 was obtained from Merck (Darmstadt, Germany).

### Organisms, media, and culture conditions

The ascomycete *Fusarium equiseti* K3 was isolated from HCH-contaminated soil collected from a former obsolete pesticide store in Kitengela, Kenya (GPS: 01.49 S, 37.048E) and its draft genome sequence was described earlier (36). The fungus was not found to degrade HCH (see below). Fungal pre-cultures were cultivated at 25°C on potato dextrose agar (PDA; Difco) prior to the transfer to ^13^C glucose/glucose-containing MSM agar. The HCH degrading bacterium *Sphingobium* sp. S8 was isolated from the same soil as the fungus and its draft genome sequence was described by Khan et al (37). The bacterium was cultivated on Luria-Bertani (LB) (DSMZ) agar medium treated with γ-HCH (97%; Sigma) to a final concentration of 100 μg/ml. To prepare the inoculum of the S8 strain, a 72-h old culture (OD_600_ ~1.5) of strain S8 in LB medium was centrifuged at 12 000 g at 15°C for 10 min, the cell pellet washed twice with MSM medium, and resuspended in the same medium to a concentration of 7.2 ×10^8^ cells mL^−1^.

#### Inhibition test

The absence of mutual fungal-bacterial inhibition was assessed using a standard procedure as described before (38). Briefly, a fungal mycelium was inoculated in the middle of a PDA plate, and four spots of bacterial suspension (10 μL) inoculated onto the agar surface around the fungal inoculum at a distance of 1cm. The plates were later incubated at 25°C and mutual growth or growth inhibition was analyzed by visual inspection and microscopy.

### Microcosm setup

Laboratory microcosms consisted of spatially separated nutrient-rich (NR) and nutrient-deficient (ND) zones that were separated by 2 mm air gaps (**Fig. 1A**). To create NR zones, two patches (Ø 1 cm) were cut from a minimum salt medium (MSM) (39) agar plate supplemented with 10% w/v ^13^C glucose/glucose as sole carbon source as described below. The two patches (patches 1 & 3; **Fig. 1A**) were placed in a Petri dish (Ø 8.5 cm) at a distance of 1.4 cm apart. To represent a ND zone an MSM agar patch (l x w x h: 1 ×1 × 0.5 cm; patch 2) was placed between them, leaving a 2 mm air gap on either side (**Fig 1A**). Patch 2 served as a spot for HCH spiking (in the case of HCH degradation setups) and bacterial inoculation. Fungal strain K3 was inoculated on patch 1 and allowed to grow until its hyphae touched the edges of patch 2 (after ca. 3 d). Fifty μL (i.e., ≈ 3.6 × 10^7^ cells) HCH degrading strain S8 in MSM medium was applied to patch 2 of the microcosm. The microcosms were then incubated at 25 °C in the dark until the mycelia had overgrown the patches (ca. 10 d) and subsequently subjected to protein and HCH extraction and analysis.

**Figure 1:**
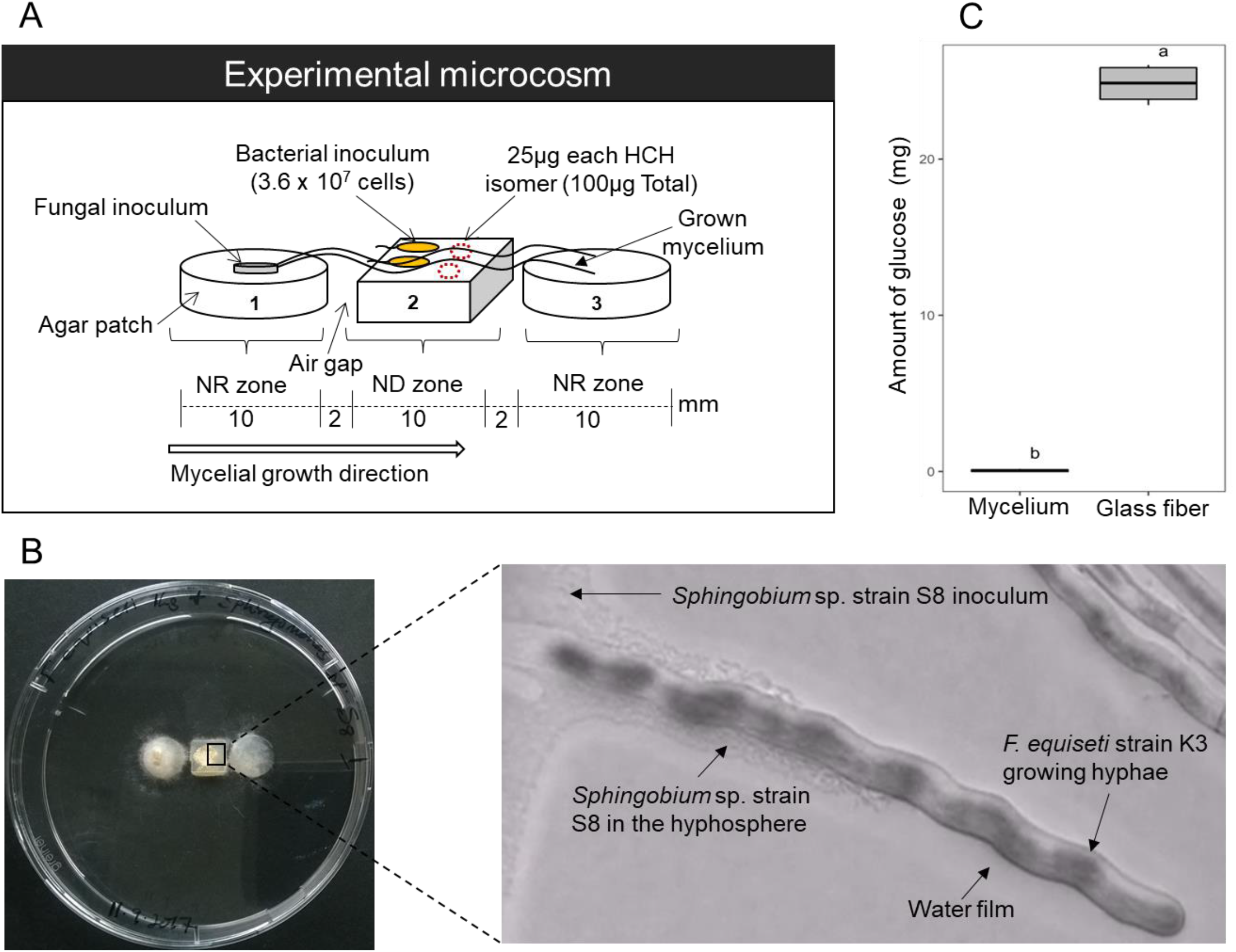
Fragmented synthetic microcosms to assess mycelial nutrient transfer and its effect on co-metabolic bacterial HCH degradation. (A) Scheme of the microcosm consisting of two nutrient rich (NR) agar patches (patch1 and patch 3) and a nutrient-deficient agar patch 2 (ND zone) separated by air gaps to mimic unsaturated soil habitats. The fungus was inoculated to patch 1 and allowed to overgrow bacteria on patch 2 in presence of HCH. (B) Bird’s view of the setup placed in a Petri dish and micrographs of *Sphingobium* sp. S8 growing as biofilm in close proximity to growing hyphae of *F. equiseti* K3 on patch 2. (C) Glucose content measured on patch 2 after 10 days in the presence of *F. equiseti* K3 and glass fiber. Different lowercase letters indicate statistically significant difference between the treatment groups (p < 0.05, *n* = 3).

### Analysis of fungal nutrient transfer to bacteria

#### Preparation of microcosms and inoculation

Patch 1 (**Fig. 1A**) was spiked with ^13^C glucose by liquefying MSM medium agar at 95°C in a water bath and subsequently transferring a five mL aliquot into a sterile tube containing 0.5 g of ^13^C glucose. The tube was then warmed at constant shaking at 750 rpm in a thermomixer (Eppendorf, Germany) at 95°C until complete glucose dissolution. Two droplets of 200 μL each of the ^13^C-glucose-agar (i.e., 40 mg glucose) were then placed into the Petri dishes (Ø 8.5 cm) at a distance of 1.4 cm apart and allowed to solidify under a sterile laminar flow hood to form patches 1 & 3) (Fig. 1A) while patch 2 was used an inoculation spot for the HCH degrading bacteria. Parallel control setups including glass fiber experiments to mimic mycelial structures and control experiments with bacteria directly growing on a ^13^C-glucose enriched (~20 mg) agar patch were prepared (**Fig S1.1**). Following a 10d incubation period, the microcosms were subjected to protein and glucose extraction from patch 2 and analyzed as described below. All experiments were performed in triplicate. Identical setups in the absence of bacteria served to quantify the glucose amounts transported by diffusion or capillary forces along glass fibers from patch 1 to patch 2 in the presence and absence of glass fibers after 10d. To do so, patch 2 was weighed and transferred into a 15 mL Falcon tube (Corning, USA). Subsequently, 5 mL of hot water and 3 mL of 1N HCl were added and the tube vigorously shaken for 5 minutes. The suspension was then filtered and the glucose concentration measured by pulsed amperometric detection (PAD).

#### Protein extraction, sample preparation

After harvest, patch 2 (**Fig 1A**) was transferred to sterile 10 mL glass tubes containing 2 mL of 1x PBS buffer pH 7.4, the tubes vortexed for 1 min and then sonicated (3 times for 30 s with 1 min pause each time) in a sonication bath (SONOREX SUPER RK 255 H, BANDELIN, Germany) at 35kHz ultrasound wave. The cell suspension was then transferred into 2 mL tubes and centrifuged at 16 000g and 4°C (Eppendorf, Germany) to pellet the cells. The cells were lysed in 50 μL lysis buffer (10 mM Tris; 5 mM EDTA; 0.29% NaCl; and 0.4% SDS) and processed as previously described (40). A Bradford assay was used to estimate protein concentrations. For the SDS-PAGE, 75 μg of protein was precipitated with an equal volume of 20% trichloroacetic acid (TCA, Sigma) for 16 h at 4°C. The proteins were then separated in a 12% acrylamide gel using the Hoefer^™^ mini-Vertical Electrophoresis system (Thermo Fisher Scientific, USA) and the Laemmli-buffer system (40). The protein bands were visualized by staining the gel in colloidal Coomassie Brilliant blue G-250 (Roth, Kassel, Germany). Protein bands were excised from the gel and subjected to ingel tryptic digestion as described previously (40). Tryptic peptides of all gel slices were then extracted for LC-MS/MS analysis.

#### Protein-SIP analysis and evaluation

Peptides were analyzed by LC-MS/MS as described before (32): after tryptic digestion of the samples, the resulting peptides were subjected to a shotgun proteomics workflow (nanoLC-MS/MS). Briefly, peptide lysate was injected into a nanoLC system (UltiMate 3 000, Dionex, Thermo Fisher Scientific). Peptides were first trapped for 3 min on a C18-reverse phase trapping column (Acclaim PepMap^®^ 100, 75 μm x 2 cm, particle size 3 μM, nanoViper, Thermo Fisher Scientific), followed by separation on a C18-reverse phase analytical column (Acclaim PepMap^®^ 100, 75 μm x 25 cm, particle size 3 μM, nanoViper, Thermo Fisher Scientific) with a solvent flowrate of 300 nL/min and a column temperature of 35°C. Eluting peptides were ionized by a nano ion source (Advion, TriVersa Nanomate, Ithaca, NY, USA) and analyzed at the Q Exactive HF mass spectrometer (Thermo Fisher Scientific) (cf. SI **S1.2** for detailed description). Proteome Discoverer (v1.4.1.14, Thermo Scientific) was used for protein identification and the acquired MS/MS spectra were searched with the SequestHT algorithm against the UniProt reference proteome of *Sphingobium* sp. and *Fusarium equiseti.* Trypsin served as cleavage enzyme, allowing a maximum of two miss cleavages. The precursor mass tolerance (MS) was set to 10 ppm, and the fragment mass tolerance (MS/MS) was 0.05 Da. Carbamidomethylation of cysteine was considered as fixed and oxidation of methionine was set as dynamic modification. Peptide spectrum matches (PSM’s) were validated using percolator with a false discover rate (FDR) <1% and quality filtered for XCorr ?2.25 [for charge state +2] and ≥2.5 [for charge state +3]. Identified proteins were grouped by applying the strict parsimony principle. Identifi cation of ^13^C-labeled peptides and quantification of ^13^C incorporation was done by comparing measured and expected isotopologue patterns, chromatographic retention time, and fragmentation patterns using MetaProSIP as previously described (32). Proteome data from the genomes of *Sphingobium* sp. S8 and *Fusarium equiseti* K3 were used as a decoy input file in the MetaProSIP pipeline. Data was evaluated by calculating the relative isotope abundance (RIA) and the labeling ratio (LR) of the labeled peptides.

### Analysis of co-metabolic HCH degradation

#### HCH quantification

Similar setups as described above in presence of unlabeled glucose were used to quantify the effects of mycelial C-substrate transfer on bacterial degradation of *α-, β-, γ-,* and *δ*-HCH isomers, where 10 μL of a HCH mix dissolved in acetone containing 25μg of each isomer and Fifty μL (i.e., ≈ 3.6 × 107 cells) of bacterial cells suspended in MSM media were simultaneously placed to patch 2 (**Fig 1A**). Four circular agar patches containing activated carbon (60 mg mL^−1^) were included in the microcosms to decrease the gaseous phase concentration of HCH. After a 10 d incubation period, the patches (patch 1 – 3 and the activated carbon agar patches) were separately transferred into clean 40 mL screw cap septum glass vials (Thermo Scientific, USA), crushed with 10–15 g of dried sodium sulfate, and then extracted with hexane/acetone (3:1, v/v) containing 0.31 μgL^−1^ dichlorodiphenyldichloroethylene (DDE) as an internal standard. One hundred μL of the extract was mixed with 10μL of 4-chlorobenzotrichloride as injection standard and analyzed by GC (HP 7890 Series GC, Agilent, USA) with a 20 m 0.18mm (18μm film thickness) HP5MS capillary column (Agilent, USA). Helium was used as a carrier gas at a flow rate of 1 mL min^−1^. Quadruplicate setups including an abiotic setup, a fungus-only, and a bacteria-only setup, were also tested as controls (cf. **Fig S1.2**).

#### Data analysis

To account for possible variations of the amounts of the HCH isomer added (e.g., due to volatilization, varying dissolution properties of the isomers), the amounts of the individual isomers recovered immediately after application onto patch 2 were taken as the initial amounts of HCH (*A*_t0_). Subsequently, HCH amounts (*A*_deg, t10_), and the residual (*F*_res, t10_) and degraded fractions (*F*_deg, t10_) of added HCH after 10 d were derived from equations 1–4 (cf. SI **S2.3** for details). *A*_t10_ is the residual amounts of HCH, *A*_abiotic, t10_ the residual HCH amounts in the abiotic control, *A*_p1, t10 or P2, t10 or P3,t10_ the residual HCH amounts on patches 1, 2 or 3, and *A*_AC,t10_ the residual HCH amount on activated charcoal.

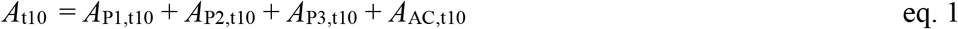

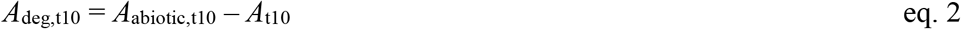

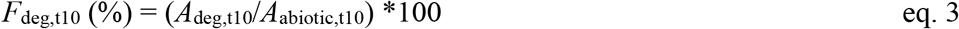

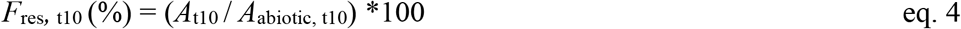

Statistical testing of the experimental treatments (abiotic, fungal-monoculture, bacterial-monoculture, and fungal-bacterial co-culture; p < 0.05) was performed on residual HCH concentrations (*A*t10) using analysis of variance (one-way ANOVA) and Tukey’s HSD (honestly significant difference) in R (v 4.2.0). The significance of differences between individual means was computed using the student’s t-test.

## Results

### Uptake of hyphal nutrients analyzed by protein-SIP

In a spatially heterogeneous laboratory ecosystem, we assessed whether mycelial nutrient transfer from nutrient-rich to nutrient-deprived areas facilitates bacterial co-metabolic HCH degradation in a nutrient-deficient habitat. Using ^13^C-labelled fungal biomass and protein-SIP, we observed higher protein contents and stable isotope enriched proteins in *Sphingobium* sp. S8 in presence of fungal hyphae (protein content: 0.45 mg) as compared to controls in presence of glass fiber (0.37 mg) and the absence of hyphal or glass fiber transport (0.29 mg; SI **Table S2.1**). The relative isotope abundance (RIA, **Fig. 2**), the labeling ratio (LR; **Fig. 2**) and the shape of isotopic spectral patterns (**Fig. 3**) evidenced ^13^C enrichment of the total bacterial proteome (RIA = 10.9 – 93.9%; LR = 0.15 – 0.57) and selected proteins (RIA = 10.1 – 92%; LR = 0.17 – 0.59) (SI **Table S2.2**). Constitutively expressed bacterial proteins from core cellular and metabolic processes were selected out of 87 labeled proteins for assessing ^13^C incorporation in our treatments (**Fig. 2** and **Table S2.2**): the chaperone DnaK (RIA = 9.4%; LR = 0.13), the ATP synthase subunit-β (RIA = 10.4%; LR = 0.24), and the peptidoglycan-associated lipoprotein (RIA = 14.2%; LR = 0.14). To quantify fungal impacts on the expression of central enzymes of bacterial HCH degradation, the ^13^C incorporation into two enzymes responsible for the initial steps of HCH degradation was quantified (Figs. 2&3); i.e. the haloalkane dehalogenase (LinB; RIA = 9.5%; LR = 0.35) and the 2,5-dichloro-2,5-cyclohexadiene-1,4-diol dehydrogenase (Lin C; RIA = 7.1%; LR = 0.34). LinB and LinC exhibited significant labeling and hence were built on ^13^C enriched substrates upon hyphal transfer. As expected, observed RIA (88% – 96%) and LR (0.38 - 0.82) values in hyphal nutrient transfer experiments were different (p <9.8 × 10^−5^) in cells cultivated on ^13^C glucose agar. Control experiments applying glass fibers to link patches 1 and 2 instead of mycelia were further performed assuming that abiotic fibers may not actively transfer ^13^C glucose and, hence, would not lead to increased RIA and LR. Likely due to capillary effects, efficient glucose transport to patch 2 was observed leading to clear glucose enrichment (≈24.8 mg) as found in additional abiotic glucose transport experiments (**Fig. 1C)**. By contrast, no hyphal glucose transfer to bacteria-free patch 2 (< 0.14 mg) was measured (**Fig. 1C**), hence suggesting that ^13^C enrichment of the proteins in the fungal transfer experiments was due to fungal metabolites. Such assumption is further backed by RIA and LR values of glass fiber experiments, where RIA (86% – 97%) was similar to bacterial cultivation experiments on^13^C glucose agar. As RIA reflects the extent to which a labeled substrate is used for biomass production (32), high RIA values in glass fiber transfer and ^13^C glucose growth experiment reflect direct assimilation of ^13^C glucose by the bacteria (41). Lower RIA values, however, were observed in presence of mycelia as a likely result of the assimilation of partially labeled fungal metabolites (32,41). The LR in the mycelial transfer experiments were similar or lower than LR in the glass fiber transfer experiments (LR = 0.15 – 0.34; **Fig 2**) yet clearly lower than on ^13^C glucose agar (LR = 0.38 – 0.82). Peptide mass spectra patterns also reflected ^13^C protein incorporation (**Fig 3**) and allowed for the visual distinction between direct assimilation of the labeled substrate (**Fig 3B, C**) or utilization from partially labeled metabolites (**Fig 3A**) secreted by other organisms (32). The skewed isotopic incorporation pattern (**Fig 3A**) in presence of mycelia, therefore, suggests assimilation of partially labeled substrates (41).

**Figure 2:**
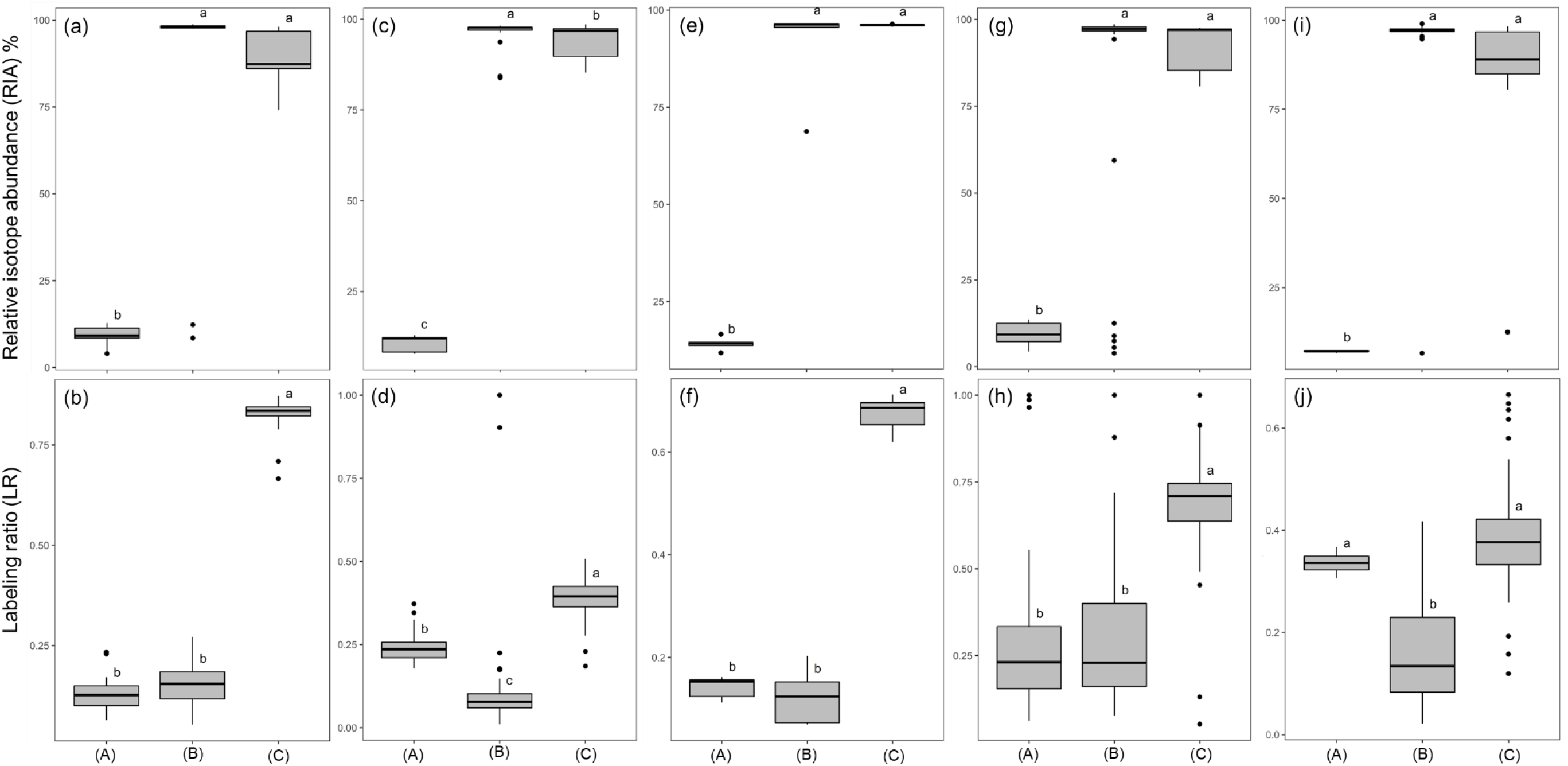
Relative isotope abundance (RIA, Figs. 2a, c, e, g, i) and labeling ratio (LR, Figs. 2b, d, f, h, j) of housekeeping and key HCH degrading proteins of *Sphingobium* sp. S8 in presence of mycelia (A), glass fiber (B), and bacteria grown on 10% w/v ^13^C glucose agar (C). **Figs. 2 a,b**: Chaperone DnaK protein, **Figs. 2c&d**: ATP synthase *β*-subunit, Figs. **2e&f**: peptidoglycan associated lipoprotein, **Figs. 2g & h:** haloalkane dehalogenase (LinB), and **Figs. 2i & j**: dichloro-2,5-cyclohexadiene-1,4-diol dehydrogenase (LinC) respectively. Different lowercase letters indicate statistically significant difference between the treatment groups (p < 0.05, *n* = 3).

**Figure 3:**
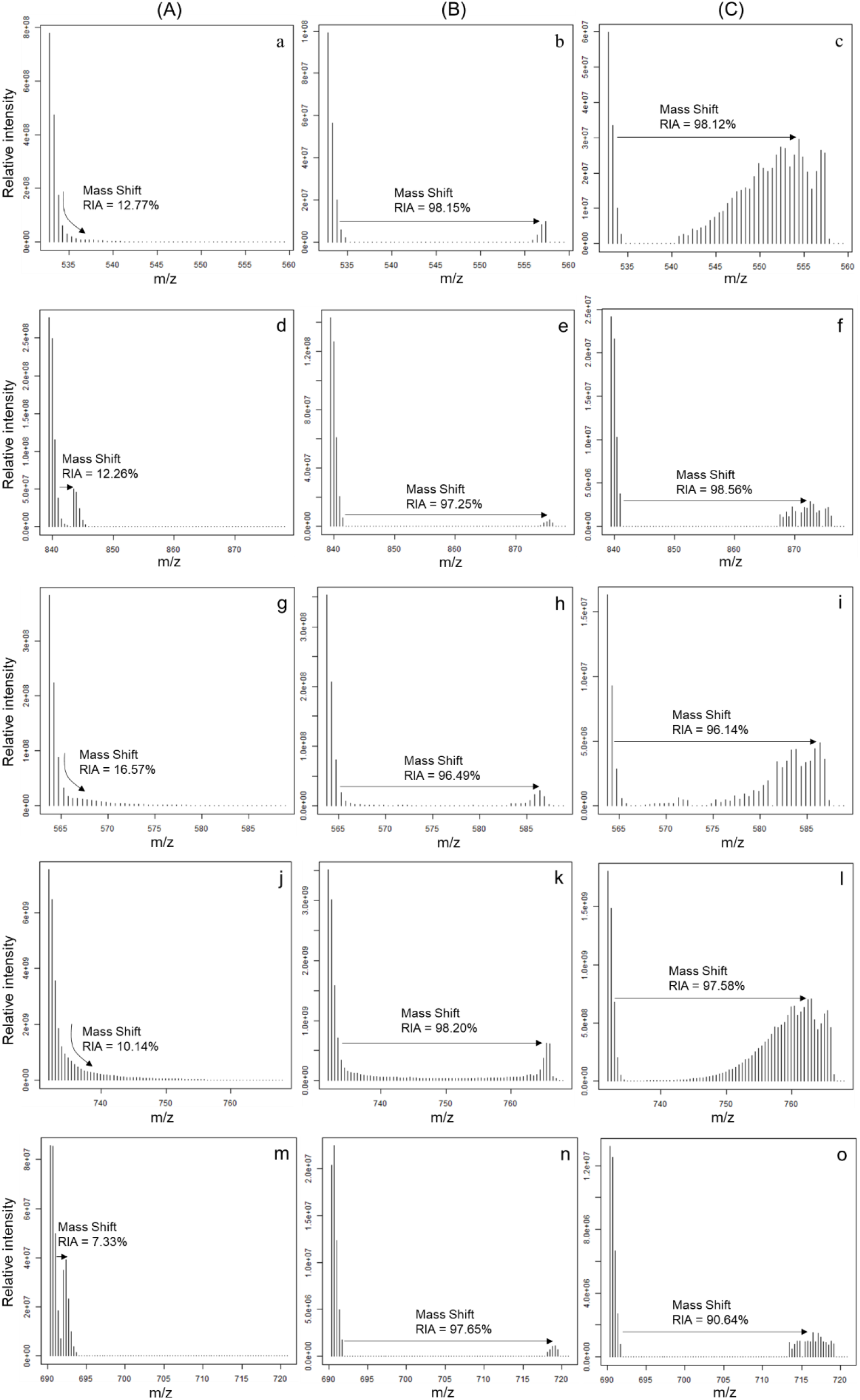
Representative mass spectra showing ion mass distribution of peptides from selected proteins on setups in presence of mycelia (A), glass fiber (B), and bacteria grown on 10% w/v ^13^C glucose agar (C) bacteria. **Figs. 3a, b, c**: peptide sequence TTPSIVAFTK of Chaperone DnaK protein; **Figs. 3d, e, f**: peptide sequence LVLEVAQHLGENTVR of ATP synthase β subunit; **Figs. 3g, h, i**: peptide sequence VTIEGHADER of peptidoglycan associated lipoprotein; **Figs. 3j, k, l**: peptide sequence DYAGWLSESPIPK of haloalkane dehalogenase; **Figs. 3m, n, o**: peptide sequence RGANVVVADIVEAAAEETVR of 2,5-dichloro-2,5-cyclohexadiene-1,4-diol dehydrogenase.

### HCH biodegradation by *Sphingobium* sp. S8

We further quantified the effects of mycelial C-substrate transfer on bacterial HCH degradation. Lower residual HCH fractions (*F*_res, t10_; eq. 4) of all HCH isomers were found in fungal-bacterial co-cultures (*F*_res, t10_ = 19% – 45%) than in bacterial (*F*_res, t10_ = 50% – 72%) or fungal (*F*_res, t10_ = 92% – 99%) mono-cultures after 10 days of incubation (SI **Table S2.4)**. No difference in *F*_res, t10_ (p < 0.05) was observed between the fungal monoculture and abiotic control setups (SI, **Fig S2.1**) suggesting that increased HCH degradation in presence of fungal-bacterial co-cultures was due to increased bacterial activity. To distinguish between abiotic and biotic losses, the HCH fractions lost by microbial activity (*F*_deg, t10_) were calculated (eq 3) revealing that 71 % (*α*-HCH), 57 %, (*β*-HCH), 75% (*γ*-HCH), and 81% (*δ*-HCH) were degraded by fungal-bacterial activity (**Fig. 4**). For the *α-, β-, γ-,* and *δ*-HCH isomers this results in ca. 1.9-, 1.1-, 1.6- and 1.6-fold better degradation than by a bacterial monoculture (**Fig 4, Table S2.6**) and reveals significant (p < 0.05) influence of C-substrate transfer on the degradation of the *α*-, *γ-,* and *δ*-HCH, yet not of *β*-HCH. Batch biodegradation experiments performed in parallel revealed that *Sphingobium* sp. S8 co-metabolically degraded *β*-HCH less efficiently (~59% of 34.4 μM *β*-HCH in 24 h; SI **S1.4 & Fig S2.2**) than the other isomers (~100 % removal; SI **S1.4 & Fig S2.2**). The low degradation efficiency of *β*-HCH that is independent of additional substrates may hence explain the poor effect of hyphal transfer on the β-HCH biodegradation.

**Figure 4:**
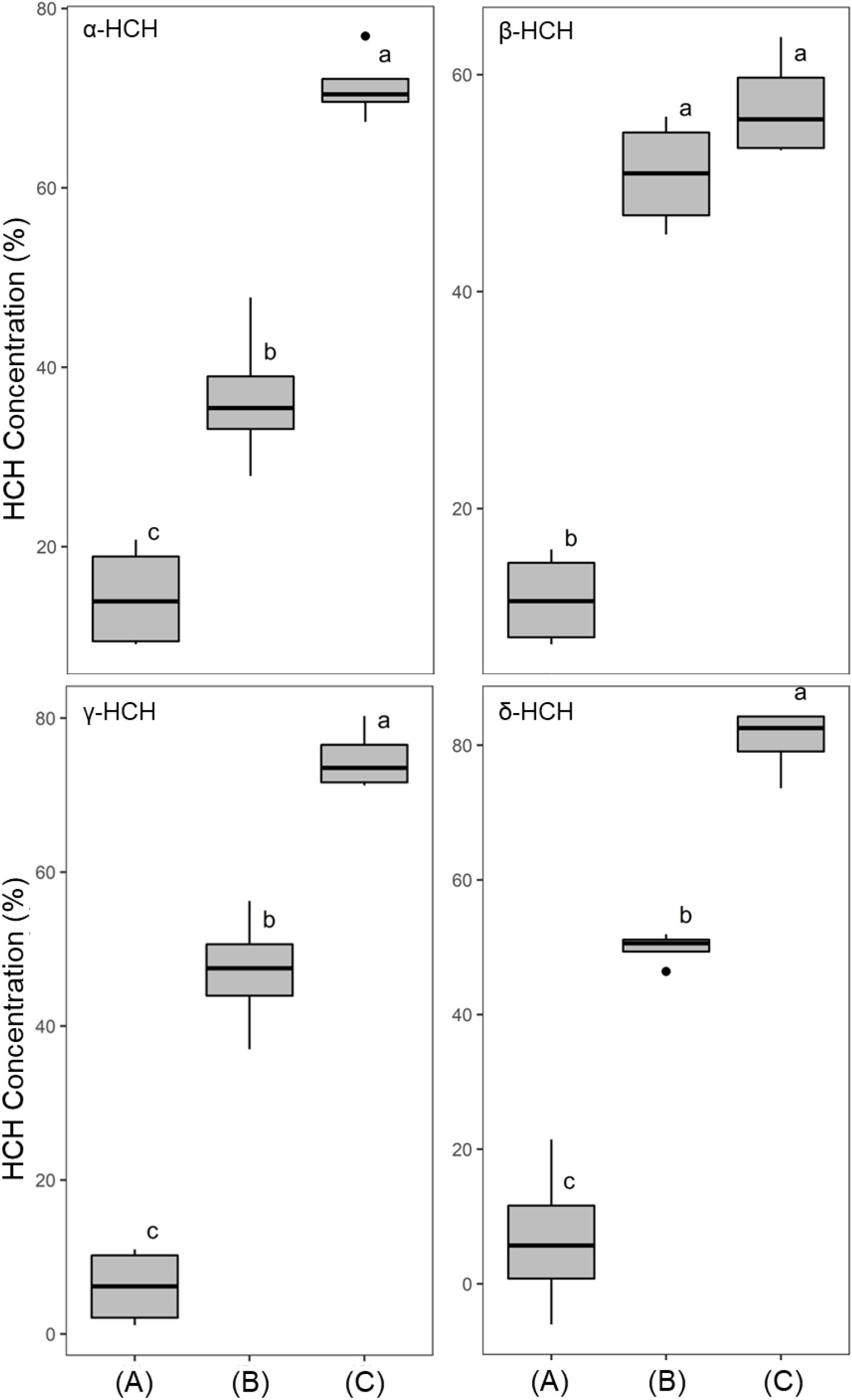
Fractions of HCH isomers degraded after 10 days of incubation (F_deg, t10_) in presence of a fungal mono-culture (B), a bacterial mono-culture (C), and fungal-bacterial co-culture (D). Different lowercase letters indicate statistically significant difference between the treatment groups (p < 0.05, *n* = 4).

## Discussion

### Hyphal transport and bacterial uptake of fungal metabolites

Using ^13^C-labelled fungal biomass and protein-SIP, we traced the incorporation of ^13^C fungal metabolites into proteins of HCH degrading *Sphingobium* sp. S8. Our findings suggest that mycelial metabolite transfer to nutrient-deficient habitats promoted bacterial growth and co-metabolic HCH in a spatially structured model ecosystem. The ^13^C-metabolite transfer was detected by relative isotope abundance, the labeling ratio, and the shape of isotopic mass distribution profiles of the bacterial proteins. Mycelial transport typically exhibited lower RIA than in fungus-free controls that provided ^13^C glucose at different bioavailability **(Fig. 1**), either by glass fiber transfer or upon growth on ^13^C glucose agar (**Figs. 2a, c, e, g, i**). The LR upon metabolite transport were similar or even higher (Chaperone DnaK (**Figs. 2 d**) and LinB (**Fig. 2j**)) than in glass fiber experiments. As the LR reflects the extent of protein turnover and hence bacterial growth (32), suggests efficient activation of the bacterial metabolism and HCH biotransformation by fungal metabolites. ^13^C incorporation into key enzymes of co-metabolic HCH degradation (i.e., LinB and LinC) thereby provides a direct link between hyphal metabolite transport and co-metabolic bacterial HCH degradation in fragmented habitats (**Fig. 4**). Our study hence provides for the first time direct molecular evidence for the role of fungal-bacterial interactions for co-metabolic contaminant degradation in an otherwise nutrient-deprived environment. In natural environments fungal-bacterial associations (18,22) can develop neutral, competitive (antagonistic) or cooperative (22,42) relationships. In a synergistic relationship as in our experiment, fungal compounds have been found to stimulate bacterial growth in the hyphosphere (14,17,22,43) by providing low molecular weight organic acids (LMWOAs) such as acetic acid and oxalic acid (44), carbohydrates, polyols, peptides, and amino acids (44,45). Although we didn’t characterize the metabolome of fungal strain K3, we speculate that similar metabolites may have led to the observed isotope HCH degradation effects. Observed active mycelial growth and the proximity of *Sphingobium* sp. S8 biofilms to fungal hyphae (**Fig 1 B**) suggests that C-substrate assimilation of strain S8 occurred by extracellular biotrophy (46) in the hyphosphere of *Fusarium equiseti* K3 as has been reported earlier in other systems (18,47,48). Such an assumption is further supported by the fact that *Sphingobium* sp. S8 (37) and *Fusarium equiseti* K3 (36) had been isolated from the same organochloride pesticide-contaminated soil.

### Co-metabolic degradation of HCH isomers

In similar setups as used for protein-SIP experiments (**Fig. S1.2**), we determined mycelial transport effects on bacterial HCH degradation by quantifying residual concentrations of the *α-, β-, γ-,* and δ-isomers of HCH. After 10 days of incubation, we did not find HCH removal by *Fusarium equiseti* K3, yet clearly increased bacterial HCH degradation in presence of mycelia (**Fig. 4 & Fig S2.1**) led to 1.6 – 1.9-fold faster biodegradation of *α*-, *γ*-, and *δ*-HCH. The least benefit (1.1-fold increase) was observed for *β*-HCH. *β*-HCH is the most recalcitrant of the HCH isomers tested, due to its equatorially substituted chlorine atoms that form a barrier to the enzymatic dehydrochlorination preferring axial chlorine atoms (7,8,49,50). *Sphingobium* sp. S8 could not effectively degrade HCH isomers in absence of an additional carbon source (**Fig 4**). In the presence of 1% glucose in liquid batch experiments, however, it showed 59-100% removal of all HCH isomers (cf. SI **Fig. S2.2**) indicating its reliance on additional carbon sources for HCH degradation.

Combining protein-SIP with the HCH degradation, our study, therefore, underpins the role of fungal exudates for efficient co-metabolic bacterial HCH degradation as also had been proposed for pesticide degradation in porous media (51,52). Co-metabolic degradation is thought to be a critical process for the removal of xenobiotic compounds at both trace and bulk concentrations (6,53,54). Benimeli et al. (2007) for instance, demonstrated simultaneous consumption of glucose and *γ*-HCH by *Streptomyces* sp. M7 with increased *γ*-HCH removal at high glucose concentrations (55). Similarly, Álvarez et al. (2012) demonstrated increased *γ*-HCH removal by *Streptomyces* strains in the presence of root exudates as an additional carbon source (56). In addition, HCH degradation in Sphingomonads have mostly been conducted in the presence of varying percentages of Luria-Bertani (LB) broth or glucose as additional carbon sources (1,57).

## Conclusion

In our study, we observed commensal incorporation of fungus-derived ^13^C-labels into bacterial peptides. The presence of ^13^C labeled bacterial peptides indicated the availability of fungal exudates for bacterial uptake and hence nutrient transfer. The labeled bacterial peptides belonged to both housekeeping enzymes and enzymes relevant to HCH degradation. The labeling of bacterial haloalkane dehalogenase (LinB) and 2,5-dichloro-2,5-cyclohexadiene-1,4-diol dehydrogenase (LinC) involved in HCH degradation indicates the functional relevance of mycelial nutrient transfer to HCH degradation by bacteria. This was indicated by improved HCH degradation (p<0.05) in the presence of fungus and bacteria compared to either bacteria (up to 2 times increase) or mycelia (up to 12 times increase) alone for all four HCH isomers. Our data thereby underpin the relevance of the mycosphere as a habitat for the efficient degradation of organic contaminants. Since co-metabolic contaminant degrading microorganisms in the mycosphere may not rely on the contaminants for growth, they also may degrade chemicals at minute concentrations (‘micropollutants’) and down to proposed cleanup endpoints in the parts per trillion range (31).

## Acknowledgment

We acknowledge the Department of Molecular Systems Biology (Proteomics lab) at the Helmholtz Center for Environmental Research (UFZ) for providing their analytical facilities for this study. Funding by Deutscher Akademischer Austausch Dienst (DAAD) and funding by a grant from the International Foundation for Science (IFS grant W/5798-1) is acknowledged. The authors thank Birgit Würz for her help with GC-MS analysis. We are also grateful to Jana Reichenbach and Rita Remer for their skilled experimental help.

## Additional Information

Supplementary information

