## supplementary information for "Mycelial nutrient transfer promotes bacterial co-metabolic organochlorine pesticide degradation in nutrient-deprived environments"

**Summary:** The Supplementary information has 15 pages consisting of 4 figures and 9 tables

*Corresponding author: Lukas Y. Wick
Mailing Address:

### S1. Materials and Methods

#### S1.1 Microcosm preparation for labeling experiment

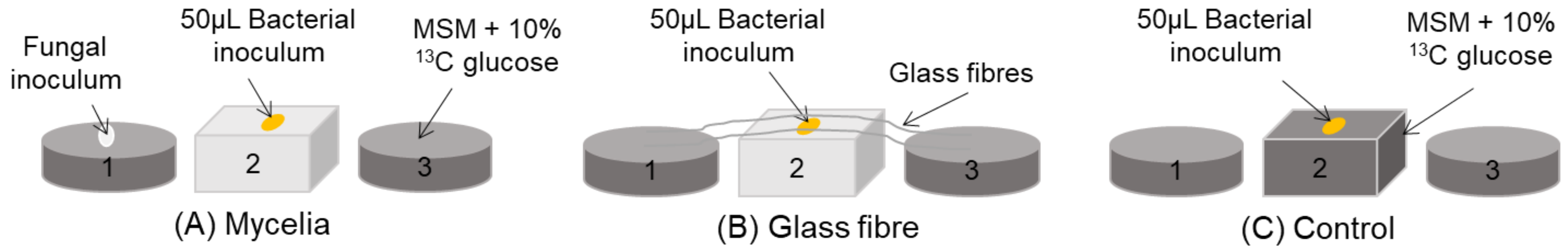


**Figure S1.1: Laboratory microcosms used for 13C labeling experiments:** (A) “mycelia”; mycelium + bacteria, (B) “glass fiber”; glass fiber + bacteria, (C) “control”; positive control used in the labeling experiment. The three agar patches were placed in a row leaving air gaps of 2 mm width in between to mimic nutrient-rich and nutrient deficient soil habitats that are separated by air-filled pores.

**S1.2 Protein-SIP analysis using nanoLC-MS/MS**

Peptides were analyzed by nanoLC-MS/MS as described before (1). After tryptic digestion of the samples, the resulting peptides were subjected to a shotgun proteomics workflow (nanoLC-MS/MS). Briefly, peptide lysate was injected into a nanoLC system (UltiMate 3000 RSLCnano, Dionex, Thermo Fisher Scientific). Peptides were first trapped for 3 min on a C18-reverse phase trapping column (Acclaim PepMap® 100, 75 µm x 2 cm, particle size 3 µM, nanoViper, Thermo Fisher Scientific), followed by separation on a C18-reverse phase analytical column (Acclaim PepMap® 100, 75 µm x 25 cm, particle size 3 µM, nanoViper, Thermo Fisher Scientific) using a two-step gradient (90 min from 4 % to 30% B, then 30 min from 30% to 55% B; A: 0.1% formic acid in MS-grade water; B: 80% acetonitrile, 0.1% formic acid in MS-grade water) with a solvent flow-rate of 300 nL $min^{-1}$ and a column temperature of 35°C. Eluting peptides were ionized by a nano ion source (Advion, TriVersa Nanomate, Ithaca, NY, USA) and analysed at the Q Exactive HF mass spectrometer (Thermo Fisher Scientific) with the following settings: MS resolution 120,000, MS automatic gain control (AGC) target 3,000,000 ions, maximum injection time for MS 80 ms, intensity threshold for MS/MS of 17,000 ions, dynamic exclusion 30 sec, TopN=20, isolation window 1.6 *m/z*, MS/MS resolution 15,000, MS/MS AGC target 50,000 ions, maximum injection time for MS/MS 120 ms.

### S1.3 Microcosm preparation for HCH degradation experiment

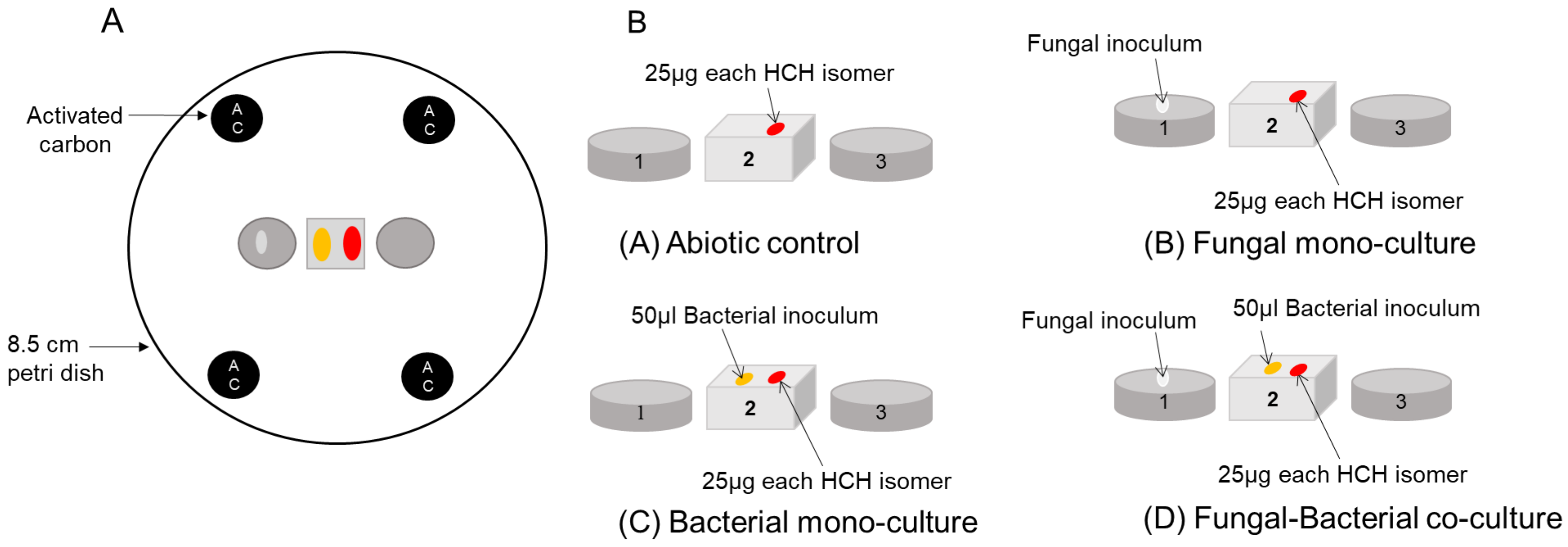


**Figure S1.2: Fragmented synthetic microcosms to quantify co-metabolic bacterial HCH degradation upon mycelial metabolite transfer.** Panel A. View of the degradation microcosm set-up from top. Panel B. Different setups used in HCH degradation experiment; (A) abiotic control (without inoculum of either the fungus or bacteria), (B) fungal mono-culture (with fungus inoculum alone), (C) bacterial monoculture (with bacterial inoculum alone), and (D) fungal-bacteria co-culture (with both fungal and bacterial inoculum). Three agar patches were placed in a row with 2mm air gaps between them to mimic nutrient-rich and nutrient-deficient contaminated pockets in soil. Agar patches containing activated carbon were included in the experiment to trap the gaseous phase HCH. All experimental setups were prepared in four replicates.

**S1.4 Biodegradation of HCH isomers by by *Sphingobium* sp. S8 in liquid-culture**

To examine HCH degradation by *Sphingobium* sp. S8, the bacterium were pre-cultured in LB medium to an $OD_{600}$ of about 1 (~3 days) and harvested by centrifugation at 7000g and 15ºC for 10min. The cells were washed twice with MSM (2) without a carbon source and resuspended in the same medium to a final $OD_{600}$ of about 3 as previously described (3). Fifty µL of cell suspension was then used to inoculate 2.5 mL of MSM medium containing 1% glucose and 10 $\mu gmL^{-1}$ (34.4µM) of each HCH isomer in a mixture in 20 ml head space GC vials that had previously been prepared. The vials were incubated in a rotary shaker at 30ºC and 150rpm. At specific time intervals (0 h, 1 h, 2 h, 5 h, 7 h, 9 h, and 24h) three vials were sacrificed for extraction and GC analysis of residual HCH in the reaction mixture. Uninoculated media were used as controls. All experiments were conducted in triplicate.

### S2. Results

#### Table S2.1 Total crude protein extracts (in mg) estimated by Bradford assay

| Experimental setup | Total crude protein extract (mg) |
|---|---|
| **Mycelia** (bacteria grown in presence of $^{13}C$ labeled mycelia) | 0.45 ± 0.06 |
| **Glass fiber** (bacteria grown in presence of glass fiber) | 0.37 ± 0.02 |
| **Control** (bacteria grown directly on $^{13}C$ glucose) | 0.66 ± 0.006 |
| **Negative control** (bacteria grown in MSM without any carbon source) | 0.29 ± 0.01 |
| ANOVA statistically significant difference [$F_{(3,8)} = 76.56$; $n = 12$, $p < 3.12 \times 10^{-6}$] | |

#### Table S2.2 Computed average RIA and LR of the total bacterial proteome and selected proteins from different experimental setups

| | RIA (%) | | | LR | | |
|---|---|---|---|---|---|---|
| **Proteins** | **Mycelia** | **Glass fiber** | **Control** | **Mycelia** | **Glass fiber** | **Control** |
| Chaperone DnaK | 9.4 ± 2.6 | 88.8 ± 27.6 | 89.2 ± 6.9 | 0.13 ±0.04 | 0.15 ±0.05 | 0.82 ±0.05 |
| ATP synthase subunit-β | 10.4 ± 2.1 | 96.9 ± 2.5 | 94.2 ± 4.1 | 0.24 ±0.05 | 0.11 ±0.16 | 0.38 ±0.05 |
| Peptidoglycan associated lipoprotein | 14.2 ± 1.7 | 90.7 ± 12.2 | 96.2 ± 0.1 | 0.14 ±0.02 | 0.12 ±0.06 | 0.67 ±0.03 |
| Haloalkane dehalogenase (LinB) | 9.5 ± 3.0 | 86.6 ± 28.8 | 92.7 ±6 .0 | 0.35 ±0.31 | 0.34 ±0.26 | 0.67 ±0.16 |
| dichloro-2,5-cyclohexadiene-1,4-diol dehydrogenase (Lin C) | 7.06 ± 0.27 | 94.3 ± 16.0 | 87.8 ± 14.9 | 0.34 ±0.02 | 0.15 ±0.01 | 0.39 ±0.13 |
| Total selected proteins | 10.1 ± 2.6 | 91.4 ± 4.2 | 92.0 ± 3.5 | 0.24 ±0.10 | 0.17 ±0.09 | 0.59 ±0.19 |
| Total proteome | 10.9 ± 0.27 | 93.9 ± 17.0 | 92.9 ± 11.4 | 0.22 ±0.17 | 0.15 ±0.12 | 0.57 ±0.19 |

### S2.3 Estimation of HCH degradation efficiency

To account for possible variations of the amounts of the four HCH isomer added (e.g., due to volatilization of the isomers), the amounts of the individual isomers recovered immediately after application onto patch 2 were taken as the initial amounts of HCH ($A_{t0}$). Subsequently, HCH amounts ($A_{deg,\ t10}$), and the residual ($F_{res,\ t10}$) and degraded fractions ($F_{deg,\ t10}$) of added HCH after 10 d were derived from equations eqs. S1– 4. $A_{t10}$ is the residual amounts of HCH, $A_{abiotic,\ t10}$ the residual HCH amounts in the abiotic control, $A_{P1,\ t10\ or\ P2,\ t10\ or\ P3,\ t10}$ the residual HCH amounts on patches 1, 2 or 3, and $A_{AC,\ t10}$ the residual HCH amount on activated charcoal. $A_{air,\ t10}$ is the HCH amount lost by volatilization after 10 days and $F_{air,\ t10}$ (%) the corresponding fraction of HCH lost by volatilization (cf. **Tables S2.3.1- S2.3.4 & Table S2.4)-**.

$$A_{t10} = A_{P1,t10} + A_{P2,t10} + A_{P3,t10} + A_{AC,t10} \qquad \text{eq. S1}$$

$$A_{deg,\ t10} = A_{abiotic,\ t10} - A_{t10} \qquad \text{eq. S2}$$

$$F_{deg,t10}\ (\%) = (A_{deg,t10}/A_{abiotic,t10})\ {*}100 \qquad \text{eq. S3}$$

$$F_{res,\ t10}\ (\%) = (A_{t10}\ /\ A_{abiotic,\ t10})\ {*}100 \qquad \text{eq. S4}$$

$$A_{air,\ t10} = A_{abiotic,\ t10} - A_{t0} \qquad \text{eq. S5}$$

$$F_{air,\ t10}\ (\%) = (A_{air,\ t10}/A_{t0})\ {*}100 \qquad \text{eq. S6}$$

**Table S2.3.1 Estimated HCH degradation efficiency for *α*-HCH (cf. S2.3 eqs. S1 – 4)** based on residual HCH amounts after 10 days in quadruplicate laboratory microcosms (Fig. 1A) in presence of *Sphingobium* sp. strain S8 (bacterial mono-culture), *Fusarium equiseti* strain K3 (fungal mono-culture) and fungal bacterial co-cultures.

| ***α*-HCH** | | | | | | | | | |
|---|---|---|---|---|---|---|---|---|---|
| **Experimental setups** | $A_{t0}$ | $A_{P1, t10}$ | $A_{P2, t10}$ | $A_{P3, t10}$ | $A_{AC, t10}$ | $A_{t10}$ | $A_{deg, t10}$ | $F_{deg, t10}$ | $F_{air, t10}$ |
| Abiotic control 1 | 23.558 | 0.012 | 3.274 | 0.010 | 13.882 | 17.178 | | | 27.1 |
| Abiotic control 2 | 21.107 | 0.008 | 3.697 | 0.012 | 14.647 | 18.365 | | | 13.0 |
| Abiotic control 3 | 21.352 | 0.006 | 3.637 | 0.006 | 13.570 | 17.219 | | | 19.4 |
| Abiotic control 4 | 26.819 | 0.007 | 4.073 | 0.007 | 15.892 | 19.979 | | | 25.5 |
| Fungal mono-culture 1 | 23.558 | 1.006 | 0.814 | 0.214 | 12.829 | 14.864 | 3.321 | 18.3 | |
| Fungal mono-culture 2 | 21.107 | 0.776 | 0.470 | 0.230 | 14.970 | 16.446 | 1.739 | 9.6 | |
| Fungal mono-culture 3 | 21.352 | 0.942 | 0.763 | 0.182 | 14.639 | 16.527 | 1.659 | 9.1 | |
| Fungal mono-culture 4 | 26.819 | 0.484 | 0.501 | 0.041 | 13.382 | 14.407 | 3.778 | 20.8 | |
| bacterial mono-culture 1 | 23.558 | 0.056 | 1.027 | 0.121 | 10.640 | 11.845 | 6.340 | 34.9 | |
| bacterial mono-culture 2 | 21.107 | 0.010 | 1.213 | 0.025 | 8.249 | 9.497 | 8.688 | 47.8 | |
| bacterial mono-culture 3 | 21.352 | 0.009 | 1.286 | 0.032 | 10.300 | 11.627 | 6.558 | 36.1 | |
| bacterial mono-culture 4 | 26.819 | 0.025 | 1.064 | 0.016 | 12.011 | 13.115 | 5.070 | 27.9 | |
| Fungal-bacterial co-culture 1 | 23.558 | 0.160 | 0.119 | 0.020 | 3.900 | 4.199 | 13.986 | 76.9 | |
| Fungal-bacterial co-culture 2 | 21.107 | 0.342 | 0.383 | 0.015 | 4.616 | 5.356 | 12.829 | 70.5 | |
| Fungal-bacterial co-culture 3 | 21.352 | 0.200 | 0.138 | 0.008 | 5.050 | 5.395 | 12.790 | 70.3 | |
| Fungal-bacterial co-culture 4 | 26.819 | 0.153 | 0.126 | 0.038 | 5.618 | 5.936 | 12.250 | 67.4 | |

**Table S2.3.2 Estimated HCH degradation efficiency for *β*-HCH (cf. S2.3 eqs. S1 – 4)** based on residual HCH amounts after 10 days in quadruplicate laboratory microcosms (Fig. 1A) in presence of *Sphingobium* sp. strain S8 (bacterial mono-culture), *Fusarium equiseti* strain K3 (fungal mono-culture) and fungal bacterial co-cultures.

| ***β*-HCH** | | | | | | | | | |
|---|---|---|---|---|---|---|---|---|---|
| **Experimental setups** | $A_{t0}$ | $A_{P1, t10}$ | $A_{P2, t10}$ | $A_{P3, t10}$ | $A_{AC, t10}$ | $A_{t10}$ | $A_{deg, t10}$ | $F_{deg, t10}$ | $F_{air, t10}$ |
| Abiotic control 1 | 28.935 | 0.212 | 26.003 | 0.217 | 0.198 | 26.630 | | | 8.0 |
| Abiotic control 2 | 28.365 | 0.194 | 25.453 | 0.171 | 0.193 | 26.011 | | | 8.3 |
| Abiotic control 3 | 34.924 | 0.157 | 30.255 | 0.185 | 0.214 | 30.811 | | | 11.8 |
| Abiotic control 4 | 31.447 | 0.239 | 27.249 | 0.219 | 0.250 | 27.957 | | | 11.1 |
| Fungal mono-culture 1 | 28.935 | 0.565 | 24.906 | 0.250 | 0.046 | 25.767 | 2.086 | 7.5 | |
| Fungal mono-culture 2 | 28.365 | 0.521 | 24.704 | 0.270 | 0.032 | 25.526 | 2.326 | 8.4 | |
| Fungal mono-culture 3 | 34.924 | 0.870 | 22.541 | 0.309 | 0.071 | 23.791 | 4.061 | 14.6 | |
| Fungal mono-culture 4 | 31.447 | 0.641 | 22.505 | 0.127 | 0.057 | 23.330 | 4.523 | 16.2 | |
| bacterial mono-culture 1 | 28.935 | 0.595 | 11.971 | 0.184 | 0.010 | 12.760 | 15.092 | 54.2 | |
| bacterial mono-culture 2 | 28.365 | 0.108 | 14.280 | 0.182 | 0.017 | 14.587 | 13.266 | 47.6 | |
| bacterial mono-culture 3 | 34.924 | 0.075 | 14.989 | 0.168 | 0.014 | 15.245 | 12.607 | 45.3 | |
| bacterial mono-culture 4 | 31.447 | 0.216 | 11.844 | 0.147 | 0.014 | 12.221 | 15.631 | 56.1 | |
| Fungal-bacterial co-culture 1 | 28.935 | 0.130 | 9.930 | 0.101 | 0.014 | 10.175 | 17.677 | 63.5 | |
| Fungal-bacterial co-culture 2 | 28.365 | 0.133 | 12.775 | 0.068 | 0.028 | 13.005 | 14.848 | 53.3 | |
| Fungal-bacterial co-culture 3 | 34.924 | 0.071 | 12.864 | 0.114 | 0.033 | 13.082 | 14.770 | 53.0 | |
| Fungal-bacterial co-culture 4 | 31.447 | 0.108 | 11.296 | 0.133 | 0.030 | 11.566 | 16.286 | 58.5 | |

**Table S2.3.3 Estimated HCH degradation efficiency for γ-HCH (cf. S2.3 eqs. S1 – 4)** based on residual HCH amounts after 10 days in quadruplicate laboratory microcosms (Fig. 1A) in presence of *Sphingobium* sp. strain S8 (bacterial mono-culture), *Fusarium equiseti* strain K3 (fungal mono-culture) and fungal bacterial co-cultures.

| **γ-HCH** | | | | | | | | | |
|---|---|---|---|---|---|---|---|---|---|
| **Experimental setups** | $A_{t0}$ | $A_{P1, t10}$ | $A_{P2, t10}$ | $A_{P3, t10}$ | $A_{AC, t10}$ | $A_{t10}$ | $A_{deg, t10}$ | $F_{deg, t10}$ | $F_{air, t10}$ |
| Abiotic control 1 | 23.951 | 0.017 | 1.573 | 0.017 | 5.915 | 7.522 | | | 68.6 |
| Abiotic control 2 | 27.115 | 0.011 | 1.676 | 0.017 | 5.660 | 7.365 | | | 72.8 |
| Abiotic control 3 | 21.594 | 0.009 | 1.505 | 0.012 | 6.083 | 7.609 | | | 64.8 |
| Abiotic control 4 | 21.123 | 0.013 | 1.923 | 0.013 | 8.401 | 10.351 | | | 51.0 |
| Fungal mono-culture 1 | 23.951 | 1.818 | 0.391 | 0.212 | 5.589 | 8.010 | 0.201 | 2.4 | |
| Fungal mono-culture 2 | 27.115 | 1.664 | 0.229 | 0.228 | 6.186 | 8.306 | -0.095 | -1.2 | |
| Fungal mono-culture 3 | 21.594 | 1.787 | 0.362 | 0.143 | 6.735 | 9.028 | -0.817 | -9.9 | |
| Fungal mono-culture 4 | 21.123 | 1.056 | 0.242 | 0.056 | 5.955 | 7.308 | 0.903 | 11.0 | |
| bacterial mono-culture 1 | 23.951 | 0.043 | 0.440 | 0.119 | 3.606 | 4.208 | 4.004 | 48.8 | |
| bacterial mono-culture 2 | 27.115 | 0.009 | 0.509 | 0.029 | 3.046 | 3.593 | 4.618 | 56.2 | |
| bacterial mono-culture 3 | 21.594 | 0.011 | 0.590 | 0.044 | 3.767 | 4.412 | 3.799 | 46.3 | |
| bacterial mono-culture 4 | 21.123 | 0.020 | 0.487 | 0.019 | 4.650 | 5.175 | 3.036 | 37.0 | |
| Fungal-bacterial co-culture 1 | 23.951 | 0.321 | 0.038 | 0.029 | 1.232 | 1.620 | 6.591 | 80.3 | |
| Fungal-bacterial co-culture 2 | 27.115 | 0.583 | 0.130 | 0.019 | 1.628 | 2.360 | 5.851 | 71.3 | |
| Fungal-bacterial co-culture 3 | 21.594 | 0.290 | 0.047 | 0.012 | 1.682 | 2.030 | 6.182 | 75.3 | |
| Fungal-bacterial co-culture 4 | 21.123 | 0.279 | 0.061 | 0.047 | 1.929 | 2.315 | 5.896 | 71.8 | |

**Table S2.3.4 Estimated HCH degradation efficiency for *δ*-HCH (cf. S2.3 eqs. S1 – 4)** based on residual HCH amounts after 10 days in quadruplicate laboratory microcosms (Fig. 1A) in presence of *Sphingobium* sp. strain S8 (bacterial mono-culture), *Fusarium equiseti* strain K3 (fungal mono-culture) and fungal bacterial co-cultures.

| ***δ*-HCH** | | | | | | | | | |
|---|---|---|---|---|---|---|---|---|---|
| Experimental setups | $A_{t0}$ | $A_{P1,\,t10}$ | $A_{P2,\,t10}$ | $A_{P3,\,t10}$ | $A_{AC,\,t10}$ | $A_{t10}$ | $A_{deg,\,t10}$ | $F_{deg,\,t10}$ | $F_{air,\,t10}$ |
| Abiotic control 1 | 25.738 | 0.364 | 8.261 | 0.341 | 0.123 | 9.090 | | | 64.7 |
| Abiotic control 2 | 27.885 | 0.288 | 9.078 | 0.224 | 0.115 | 9.704 | | | 65.2 |
| Abiotic control 3 | 23.107 | 0.264 | 9.317 | 0.290 | 0.122 | 9.993 | | | 56.8 |
| Abiotic control 4 | 23.136 | 0.457 | 10.033 | 0.378 | 0.100 | 10.968 | | | 52.6 |
| Fungal mono-culture 1 | 25.738 | 3.616 | 3.787 | 1.648 | 0.057 | 9.108 | 0.831 | 8.4 | |
| Fungal mono-culture 2 | 27.885 | 4.309 | 4.419 | 1.771 | 0.039 | 10.537 | -0.599 | -6.0 | |
| Fungal mono-culture 3 | 23.107 | 5.068 | 3.322 | 1.162 | 0.084 | 9.636 | 0.303 | 3.0 | |
| Fungal mono-culture 4 | 23.136 | 3.571 | 3.732 | 0.440 | 0.064 | 7.807 | 2.132 | 21.5 | |
| bacterial mono-culture 1 | 25.738 | 0.190 | 4.504 | 0.075 | 0.010 | 4.779 | 5.160 | 51.9 | |
| bacterial mono-culture 2 | 27.885 | 0.068 | 4.678 | 0.172 | 0.017 | 4.936 | 5.003 | 50.3 | |
| bacterial mono-culture 3 | 23.107 | 0.085 | 5.028 | 0.199 | 0.015 | 5.327 | 4.612 | 46.4 | |
| bacterial mono-culture 4 | 23.136 | 0.105 | 4.603 | 0.157 | 0.019 | 4.884 | 5.055 | 50.9 | |
| Fungal-bacterial co-culture 1 | 25.738 | 0.322 | 1.143 | 0.097 | 0.005 | 1.567 | 8.372 | 84.2 | |
| Fungal-bacterial co-culture 2 | 27.885 | 0.377 | 1.456 | 0.061 | 0.006 | 1.900 | 8.039 | 80.9 | |
| Fungal-bacterial co-culture 3 | 23.107 | 0.216 | 1.210 | 0.122 | 0.008 | 1.556 | 8.383 | 84.3 | |
| Fungal-bacterial co-culture 4 | 23.136 | 0.326 | 2.110 | 0.178 | 0.008 | 2.622 | 7.317 | 73.6 | |

**Table S2.4: Average residual fractions of HCH isomers** based on residual HCH amounts after 10 days in quadruplicate laboratory microcosms (Fig. 1A) in presence of *Sphingobium* sp. strain S8 (bacterial mono-culture), *Fusarium equiseti* strain K3 (fungal mono-culture) and fungal bacterial co-cultures.

| $F_{res, t10}$ (%) | | | | |
|---|---|---|---|---|
| **Setups** | ***α*-HCH** | ***β*-HCH** | ***γ*-HCH** | ***δ*-HCH** |
| **Fungal mono-culture** | 97.7 ±6.8 | 91.6 ±4.6 | 99.4 ±8.7 | 93.3 ±11.5 |
| **Bacterial mono-culture** | 72.3 ±9.4 | 51.0 ±5.4 | 52.9% ±7.9% | 50.1 ±2.4 |
| **Fungal-bacterial co-culture** | 32.8 ±4.6, | 44.5 ±5.1 | 25.3 ±4.2 | 19.2 ±5.0 |

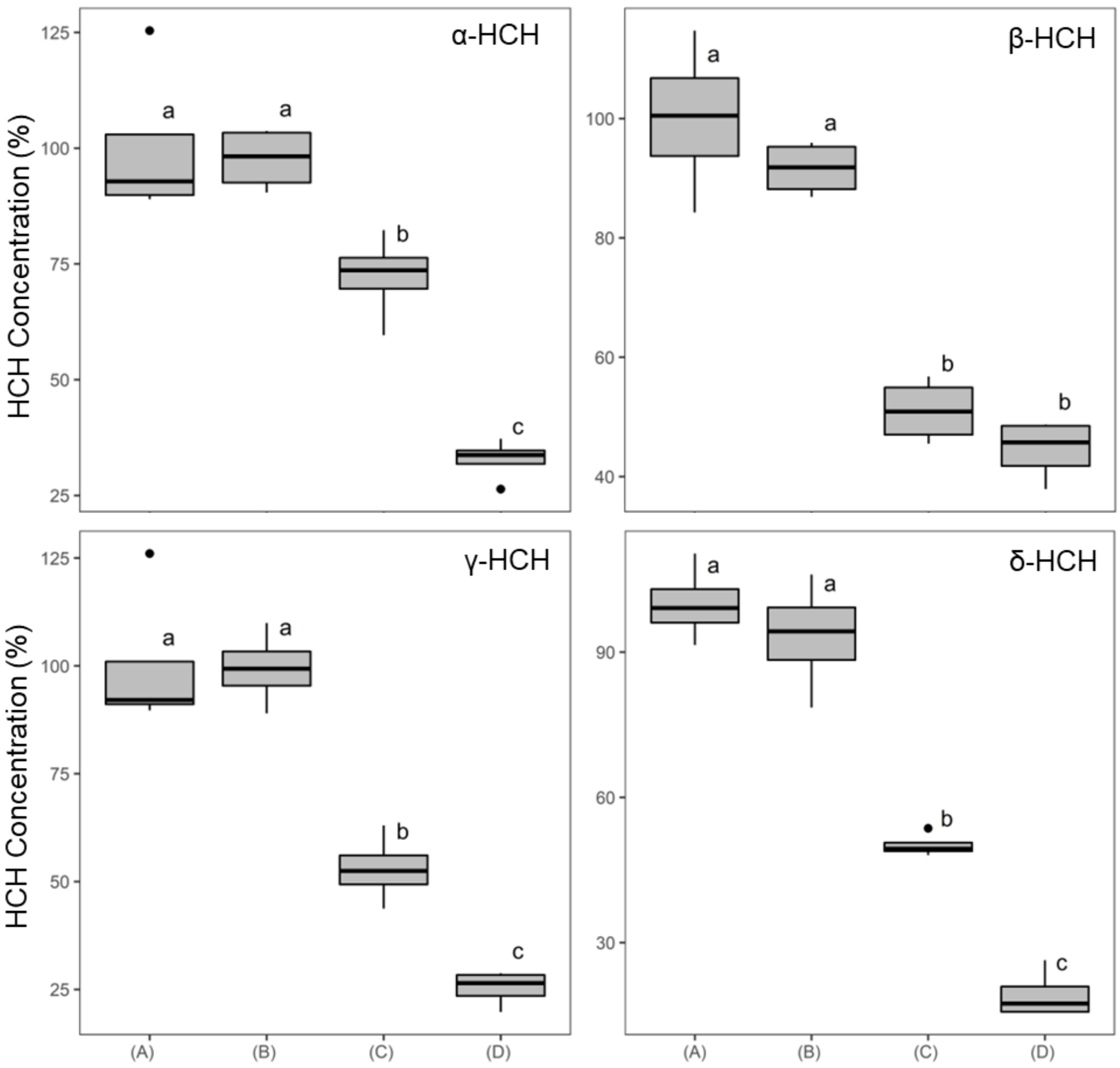


**Figure S2.1.** Residual fractions of HCH isomers after 10 days of incubation ($F_{\text{res, t10}}$) in an abiotic control (A) and in presence of a fungal mono-culture (B), a bacterial mono-culture (C), and fungal-bacterial co-culture (D). Letters a, b, and c indicate statistical significance ($p < 0.05$, $n = 4$)

**Table S2.5: Average degraded fractions of HCH isomers** based on residual HCH amounts after 10 days in quadruplicate laboratory microcosms (Fig. 1A) in presence of *Sphingobium* sp. strain S8 (bacterial mono-culture), *Fusarium equiseti* strain K3 (fungal mono-culture) and fungal bacterial co-cultures.

| $F_{deg, t10}$ (%) | | | | |
|---|---|---|---|---|
| **Setups** | **α-HCH** | **β-HCH** | **γ-HCH** | **δ-HCH** |
| **Fungal mono-culture** | 14.43 ± 5.97 | 11.66 ±4.39 | 6.14 ± 5.06 | 9.72 ± 8.11 |
| **Bacterial mono-culture** | 36.64 ± 8.25 | 50.80 ± 5.17 | 47.06 ± 7.94 | 49.88 ± 2.40 |
| **Fungal-bacterial co-culture** | 71.29 ± 4.02 | 57.07 ± 4.94 | 74.65 ± 4.15 | 80.77 ± 5.08 |

**Table S2.6: Average degradation benefits of HCH isomers in presence of fungal-bacterial co-culture as compared to fungal or bacterial mono-culture.**

| Degradation benefit | | | | |
|---|---|---|---|---|
| **Setups** | **α-HCH** | **β-HCH** | **γ-HCH** | **δ-HCH** |
| **Fungal-bacterial co-culture/Bacterial mono-culture** | 1.9 ± 0.1 | 1.1 ± 0.1 | 1.6 ± 0.1 | 1.6 ± 0.1 |
| **Fungal-bacterial co-culture/Fungal mono-culture** | 4.9 ± 0.3 | 4.9 ± 0.4 | 12.2 ± 0.7 | 8.3 ± 0.5 |

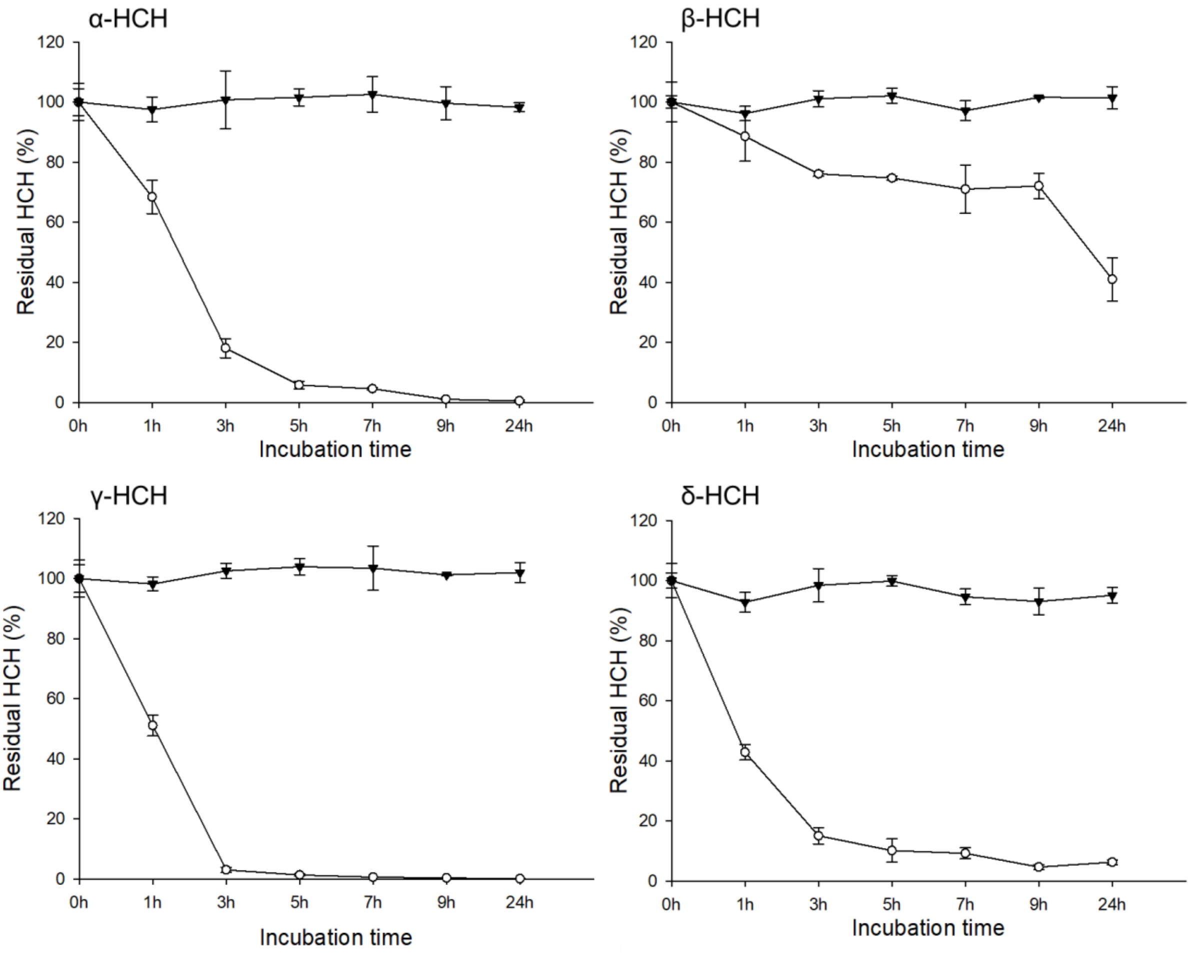


**Figure S2.2: Degradation profiles of HCH isomers by *Sphingobium* sp. S8 in liquid batch cultures.** The bacterium degraded ~98% of α-HCH, ~59% of β-HCH, ~99% of γ-HCH, and ~94% of δ-HCH, when exposed to an equal mixture of 34.4µM of each HCH isomer and 1% glucose as an additional substrate within a 24-hour incubation period.
